# Native species of the Atlantic Forest for urban environments based on functional groups: an approach to make cities in southern Brazil more resilient

**DOI:** 10.1101/2025.04.14.648749

**Authors:** Maria R Kanieski, Guilherme D Fockink, Charline Zangalli, Juliano P Gomes, Jaçanan E. de F. Milani, Cláudio Mondin, Alain Paquette

## Abstract

Planting the proper trees can mitigate and adapt cities to the adverse impacts of climate change, in particular through increasing diversity. Despite being located in one of the most biodiverse countries in the world, low diversity and overabundance of exotic species in Brazilian urban forests are common. Our objective with this study is to group species based on their functional traits to provide a tool to help planners create more resistant urban forests in cities in the domain of the Dense Ombrophilous Forest (DOF), Atlantic Forest, in southern Brazil, as well as incentivize the use of DOF species based on their functional traits. We classified 77 native trees of the DOF in Santa Catarina state, southern Brazil, with potential use in urban environments into functional groups according to the following traits: seed mass, wood density, average height, and leaf persistence, using a hierarchical cluster analysis. The different groups formed represent species with different ecological strategies. Combining species from different groups will help create more resistant urban forests in cities of the DOF domain. We hope the method will also help to increase the use of native species from DOF. Furthermore, the method can be adapted to urban forests of different domains in Brazil and other countries.

## 1. Introduction

The benefits and importance of urban forests are well-known and well-established. Urban green spaces connect people to nature and provide ecosystem services, which are of high value to communities (Zhang and Brack, 2021; Vargas-Hernandez et al., 2018). As urbanization increases, the pressure and our dependence on urban green spaces also increase, bringing several challenges to the resilience of urban trees, exacerbated by climate change (Pamuck-Albers et al., 2021; Bush and Doyon, 2019; Vargas-Hernandez et al., 2018).

Urban spaces are more vulnerable to climate change, particularly in more urbanized cities (Esperon-Rodriguez et al., 2022; Zhang and Brack, 2021). Planting the right trees can mitigate and adapt cities to the adverse impacts of climate change (Zhang and Brack, 2021; Abreu-Harbich et al., 2015), especially increasing diversity (Farrel et al., 2022; Paquette et al., 2021; Isbell et al., 2015). However, low tree diversity in urban trees is a common issue worldwide. As a result, urban greenspaces lack diversity and have low resilience to climate change because the same species will respond similarly to specific disturbances (Farrel et al., 2022; Paquette et al., 2021; Zhang and Brack, 2021).

Brazil’s urban tree canopy coverage is between 5% and 35% (mean 19%), depending on the cities (Guo et al., 2023). Furthermore, even though it is one of the most biodiverse countries in the world, it is common to see low diversity and exotic species rather than native in urban forests in Brazil (Alves et al., 2023; Soares et al., 2021). Defined as a forest of high richness, with an abundance of life forms and a complex structure, the Dense Ombrophilous Forest (DOF) of Santa Catarina, southern Brazil, in the Atlantic Forest domain, has approximately 580 species in the tree/shrub component alone, with a predominance of the Myrtaceae, Fabaceae, Lauraceae, Melastomataceae, Asteraceae and Rubiaceae families. There is also an abundance of epiphytes (Orchidaceae and Bromeliaceae) and palm trees (Lingner et al., 2013; Sevegnani et al., 2013). This high diversity of species is not reflected in the urban trees of the municipalities located within the DOF, where mostly exotic species are planted (Alves et al., 2023). The use of exotic species in urban tree planting in Brazilian cities is an old tradition passed down through generations for decades (Elias et al., 2020; Bechara et al., 2016), and reflects the lack of planning in urban trees (Núñes-Florez et al., 2019). This, however, is of course not unique to Brazil (Nock et al., 2013).

The use of native species is important to provide a specific tree identity for municipalities, connecting citizens with nature, especially because in a heavily urbanized world, people live mainly in urban ecosystems and often have little or no contact with plants that occur outside cities (Gonçalves et al., 2018; Moro and Castro, 2015). Living in an urban environment dominated by exotic species causes people to be more familiar with these species than with natives, which then impacts their own choices for planting (Moro and Castro, 2015). The presence of native species in cities could improve the knowledge of and appreciation for biodiversity, potentially increasing public support for the protection of native plants (McKinney, 2006).

The overuse of exotic species in urban forests in Brazil is also of concern for the disruption of local flora’s ecological relationships (Soares et al., 2021). The use of native species in the urban environment is preferred where possible because it contributes to the conservation of local biodiversity (Elias et al., 2020; Bechara et al., 2016). For example, they contribute to the maintenance of native fauna by providing food resources and refuge conditions, which results in more efficient dispersal and provides natural regeneration of native species (Sartori et al., 2019). Native urban forests, through nature-based solutions, play a key role in ensuring climate change mitigation and adaptation services, contributing to ecosystem services (Seddon et al., 2020) and preventing the spread of invasive species (Gonçalves et al., 2018; Moro and Castro, 2015). It should be noted, however, that exotic species are still providing important services. In some biomes with smaller pools of native species, among which few are adapted to urban life, especially in the most stressful conditions, the use of exotic species cannot be avoided. We argue that the DOF does not fit that category and has many more native species to offer than what is currently being used.

It is urgent to develop effective urban tree planning and management policies to improve the urban tree diversity that allow for the promotion and maintenance of an urban forest that is climate proof. Functional groups can be used to categorize species to help decision makers plan functionally diverse and resistant urban forests. This approach is based on fundamental principles in ecology, notably the portfolio effect (risk reduction following increased diversity) and that the ecological niche and role that species have in an ecosystem are best understood through their functional traits (Paquette et al., 2021; Yamane et al., 2018; Isbell et al., 2015; Cadotte et al., 2011; Tilman, 1999). Functional groups are simply assemblages of species that share similar traits and similar responses and sensitivity to a given stressor. The idea is, therefore, to choose species from different groups to increase functional diversity and reduce risk, much like is suggested for financial investments (simply replace species with companies and functional groups with economic sectors) (Garnier and Navas, 2012; Cadotte et al., 2011). Species from different functional groups will respond and adapt differently to unknown stressors, increasing the resistance of urban forest overall. However, their use for the planning of urban forests is still limited (Farrel et al., 2022; Paquette et al., 2021; Núñes-Florez et al., 2019). In Brazil, few studies use functional traits to evaluate the current urban forest (e. g. Pastorello et al., 2022, Monalisa-Francisco and Ramos, 2019) and to plan ahead (e. g. Zappi et al., 2022), but none are using functional groups.

Categorizing native species into functional groups allows stakeholders to identify and prioritize native species to play a key role in maintaining ecological stability and increasing biodiversity. This method not only streamlines decision-making but also highlights the unique contributions of native species to the resilience of urban ecosystems. By focusing on their functional roles, stakeholders can make a more convincing case for their protection and integration into urban environments.

This is the first study that uses functional groups as a tool for selecting native trees for urban forests in the Atlantic Forest domain. We selected 100 native trees of the Dense Ombrophilous Forest (DOF) in Santa Catarina state, southern Brazil, with potential use in urban environments. We classified the trees into functional groups according to the following traits: seed mass, wood density, average height, and leaf persistence. These traits are chosen due to their importance for plant ecological strategies (e.g. tolerance to environmental stress, dispersal mode, growth form, primary productivity, competitive ability, among others). The species selected based on the diversification of such traits helps the creation of urban forests that possess inherent resilience against a spectrum of unknown stressors, including those intensified by climate change. Our objectives were: a) to group species based on functional traits to help create urban forests that are better adapted to climate change in cities in the domain of DOF, and b) to incentivize the use of native species from DOF.

## 2. Methods

### 2.1 Study area

The area covered by this study corresponds to the region of occurrence of the Dense Ombrophilous Forest (DOF), also known as the Atlantic Forest sensu stricto, in Santa Catarina, Brazil (Fig. 1). The DOF is part of the Atlantic Forest biome, which is currently one of the world’s biodiversity hotspots, having lost approximately 93% of its original area. Although the Atlantic Forest Biome has been largely destroyed, it is still rich in biodiversity, harboring more than 8,000 endemic species of flora and fauna (Rezende et al., 2018; Myers et al., 2000). DOF is located between −25°57’40” and −29°19’13” latitude and −48°24’21” and −50°14’14” longitude. The DOF currently covers 14,907 km², corresponding to approximately 47.8% of Santa Catarina’s territory (Vibrans et al., 2021), distributed along the coastal regions from sea level to 1,000 m above sea level (Gasper et al., 2014). The climate in this forest physiognomy, according to Köppen, is classified as Cfa, humid temperate with hot summers (Alvares et al., 2013). It has high temperatures and high rainfall evenly distributed throughout the year but can also experience occasional frosts due to polar cold fronts (Oliveira-Filho et al., 2015).

**Figure 1.**
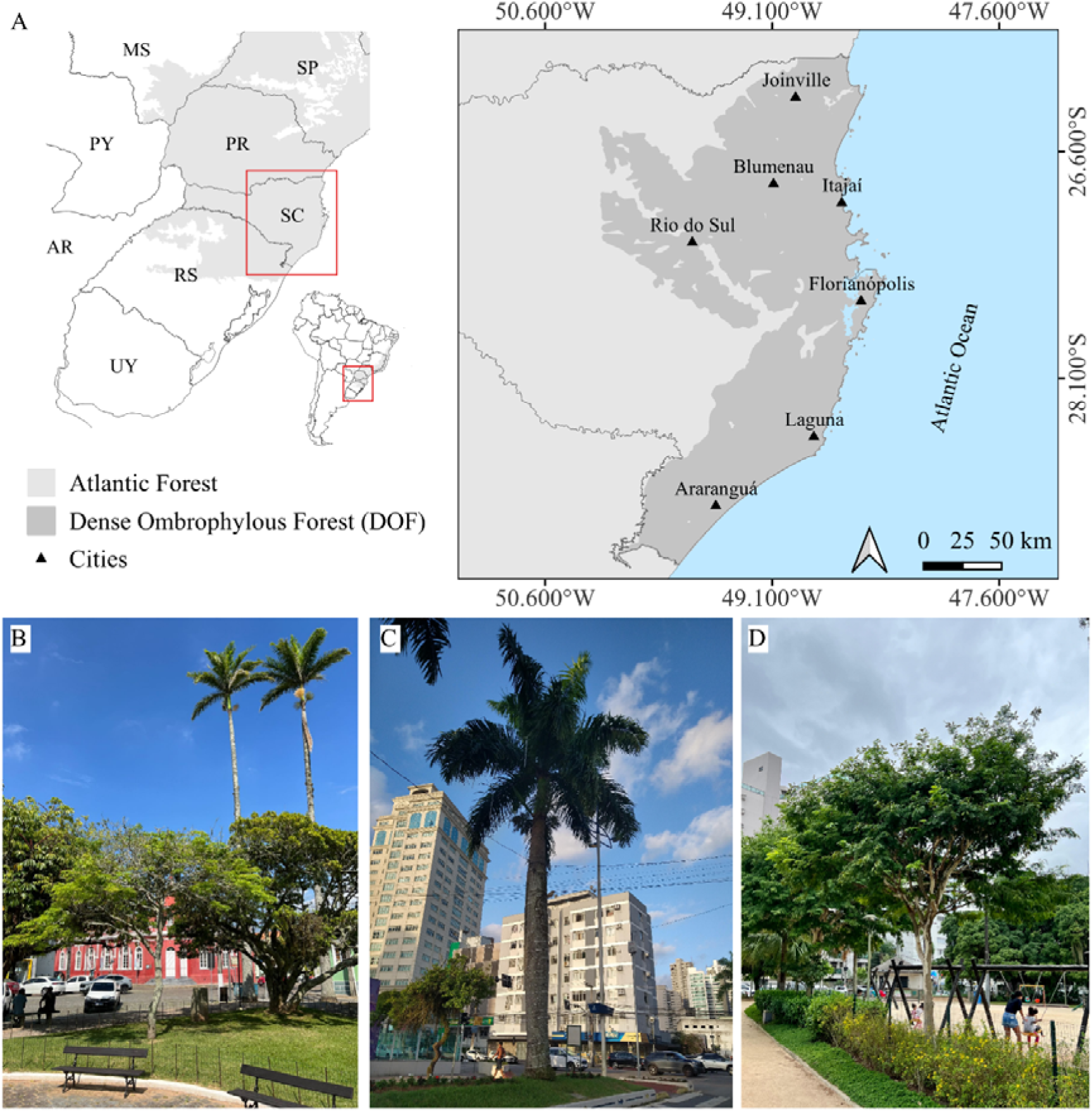
Coverage area of the Dense Ombrophylous Forest (DOF) in Santa Catarina, Brazil (A) and its main cities. Here, we present some urban spaces with trees in the cities of Laguna (B), Itajaí (C) and Florianópolis (D). We can observe the predominance of exotic species, such as A*rchontophoenix cunninghamiana* (B) and *Roystonea regia* palm trees (C) and *Libidibia ferrea* tree (D) used in the urban tree of parks and streets. Native species (B) commonly used in urban tree in these cities include *Ficus* sp. (on the right) and *Handroanthus chrysotrichus* (in the middle).

### 2.2 Species selection

We initially selected 100 native species belonging to 29 botanical families with the potential for planting in the urban forest (Table S1) from more than 500 species in the DOF in Santa Catarina (SFB, 2018). We selected the species based on relevant characteristics such as flowering (aesthetics, sent, attractivity to fauna), fruiting (attractivity to fauna, harmony with the public space), crown architecture, foliage aesthetics, and characteristics that made species compatible with the urban environment such as the absence of toxicity or aculeus, a resistant trunk and no superficial root system (USDA, 2018; Gonçalves, 2009).

### 2.3 Functional traits

Based on previous studies (Paquette et al., 2021; Núñes-Florez et al., 2019), we characterized all the species with the following functional traits: seed mass (g/seed), wood density (g/cm³), and average height (m). We chose these traits because they are commonly used in ecology due to their importance for plant ecological strategies (Wang et al., 2019; Díaz et al., 2016; Reich, 2014; Chave et al., 2009; Wright et al., 2004) and also because the trait data was available for most of the species. Seed mass is associated with dispersal mode, growth form, specific leaf area, seed dormancy, and tolerance to environmental stress, such as drought resistance, which is especially important in urban environments in the context of climate change (Harel et al., 2011; Westoby et al., 1996). Wood density is key to understanding the functioning of forest ecosystems in terms of carbon sequestration and community dynamics (Chave et al., 2005; Díaz et al., 2016) and it is an indicator of species performance (Lachenbruch and McCulloh, 2014). Plant height is a key functional trait related to aboveground biomass, leaf photosynthesis, and plant fitness, and it is a good predictor for estimating the primary productivity of terrestrial ecosystems (Díaz et al., 2016; Wang et al., 2019). In addition, we added the categorical variable leaf persistence (persistent, semi-deciduous, and deciduous), which is related to the photosynthetic rate and the competitive ability (Bernhardt-Römermann et al., 2008).

We used the three functional traits, and leaf persistence converted to factor, to assess dissimilarity between species. We obtained data through extensive bibliographic research (Flora e Funga do Brasil, 2020; Saueressig, 2014; Carvalho, 2003, 2006, 2008, 2010, 2014; Lorenzi, 1992, 1998, 2009), as well as from existing databases: Global Wood Density (Zanne et al., 2009, Chave et al., 2009), and TRY (Kattge et al., 2020).

### 2.4 Data analysis

Of the 100 species initially selected, only 77 had values for all traits and were therefore included in our analysis. We set up the groups according to the methods described in Paquette et al. (2021). In a few words, we calculated trait dissimilarity using the Gower distance for each pairwise combination of the 77 tree species in the dataset. This metric was used as it allows qualitative and quantitative variables (Pavoine et al., 2009). Subsequently, we subjected the dissimilarity matrix to a hierarchical cluster analysis based on the Ward method. We determined the cutoff point for the number of clusters by analyzing the average silhouette width followed by the biological interpretation of the clusters.

Only for the purpose of illustrating the previously computed groups and their relationships with the three continuous functional traits, we performed a principal component analysis (PCA). Leaf persistence, a categorical trait, could not be included. All analyses were performed in R version 3.5.2 (R Core Team 2021) using the agnes, daisy, and pam functions implemented in the cluster package (Maechler et al., 2023), the mutate function in the dplyr package to convert the categorical variable to factor (Wickhan et al., 2023), the PCA function from the FactoMineR package (Husson et al., 2017), the fviz_pca_bplot function from the factoextra package (Kassambara and Muntd, 2020) and the make.cepnames function from the vegan package (Oksanen et al., 2017).

## 3. Results

From the seed mass, wood density, average height, and leaf persistence traits of 77 native species from the Dense Ombrophilous Forest in Santa Catarina state, Brazil, we found six distinct functional groups (Fig. 2, Fig. 3, Table S4). The first axis of the PCA explained 42% of the variation, while the second explained 35%. Log.SM and AH showed the highest correlation with PC1, while Wd showed the highest correlation with PC2 (Fig. 4, Table S3).

**Figure 2.**
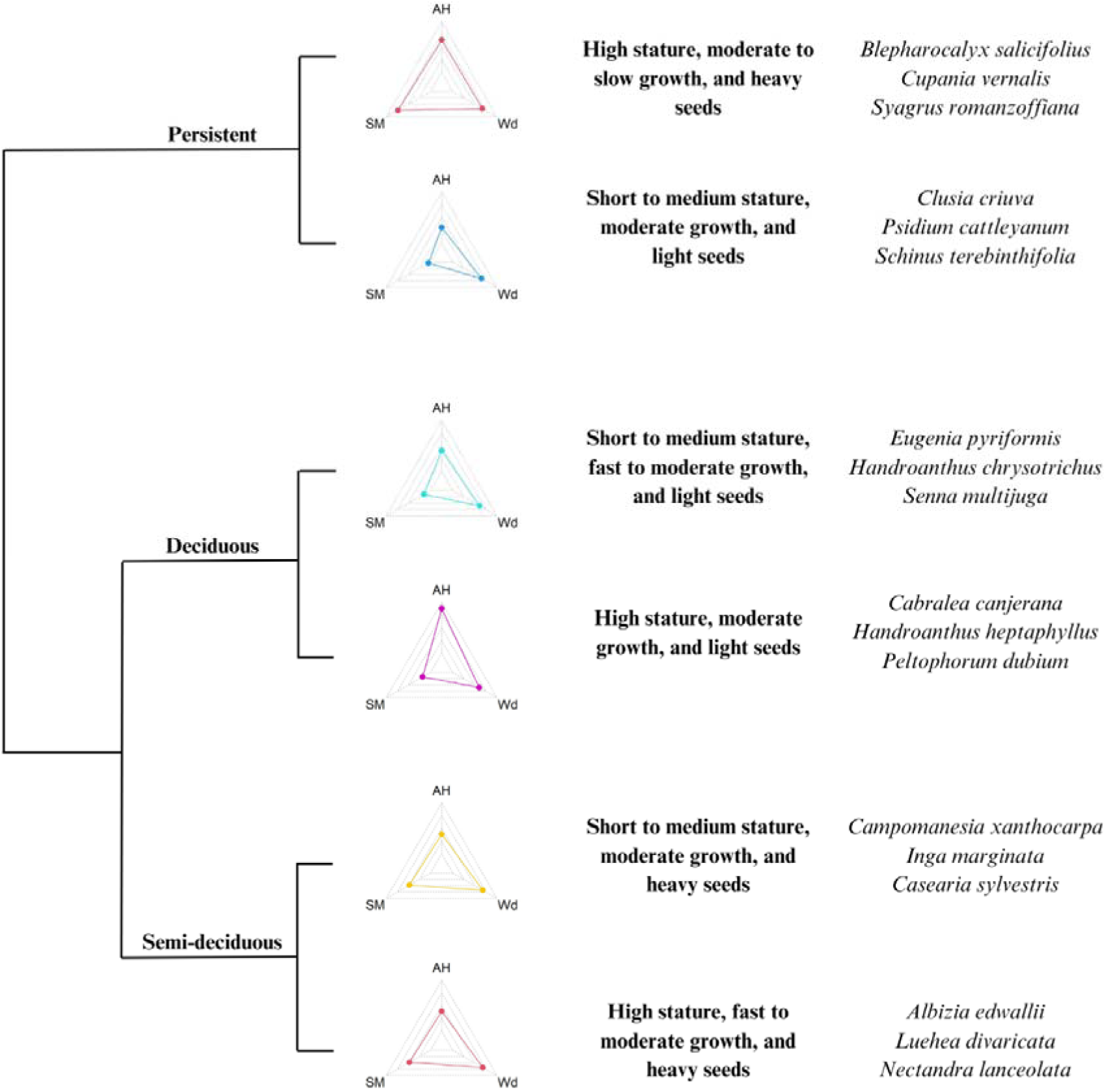
Functional groups of tree species for the Dense Ombrophylous Forest in Santa Catarina, Brazil. Functional traits: seed mass (SM), wood density (Wd), average height (AH), and leaf persistence (categorical variable; not represented) were used to construct a functional dendrogram that resulted in six distinct functional groups. Radar plots provide the relative importance of the three continuous traits in each group, pointing to different ecological strategies described in the main text, along with representative species.

**Figure 3.**
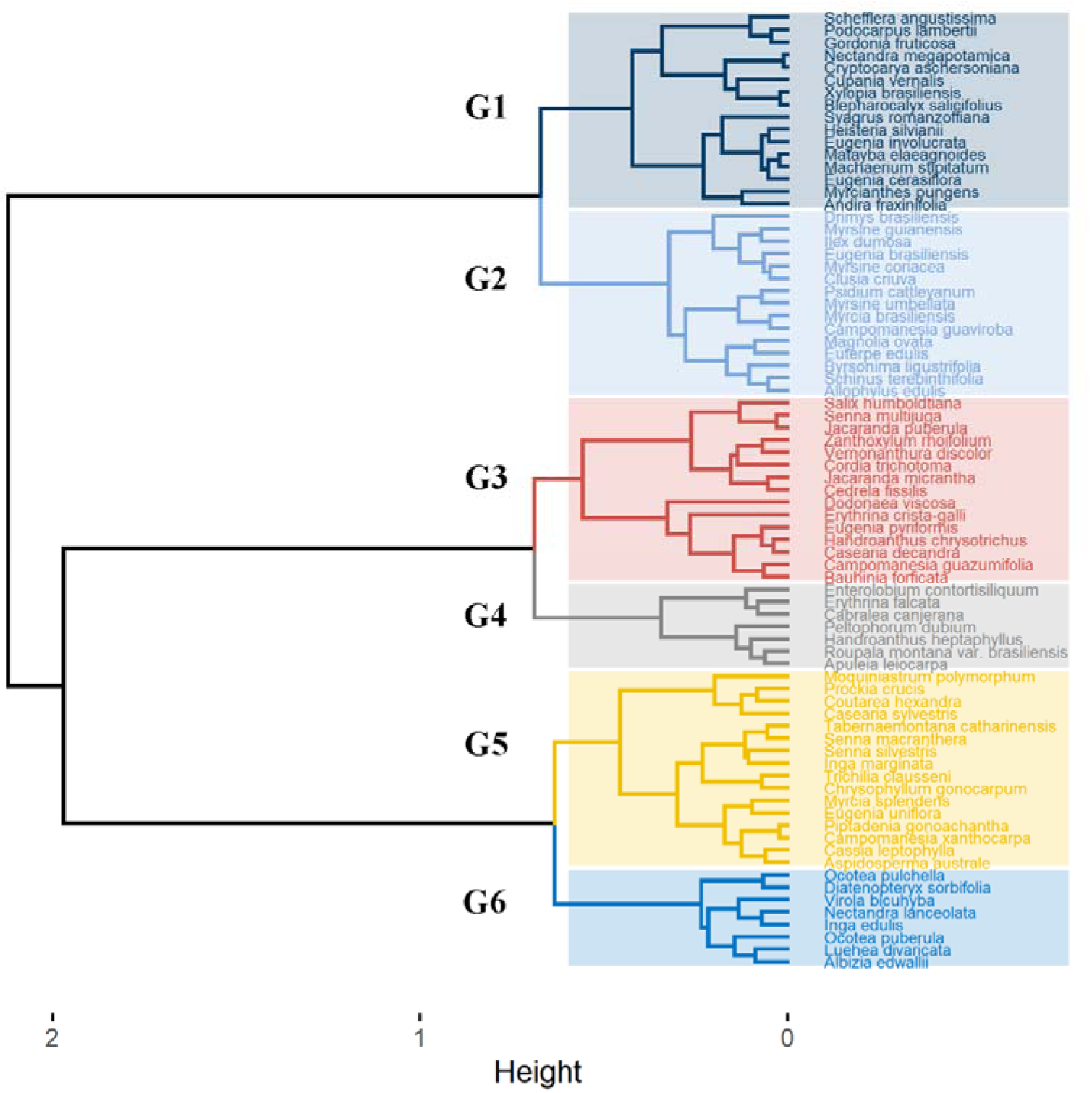
Classification of 77 native tree species of the Dense Ombrophylous Forest of Southern Brazil into six groups using a functional dendrogram, based on the similarity of their functional traits (seed mass, wood density, average height, and leaf persistence).

**Figure 4.**
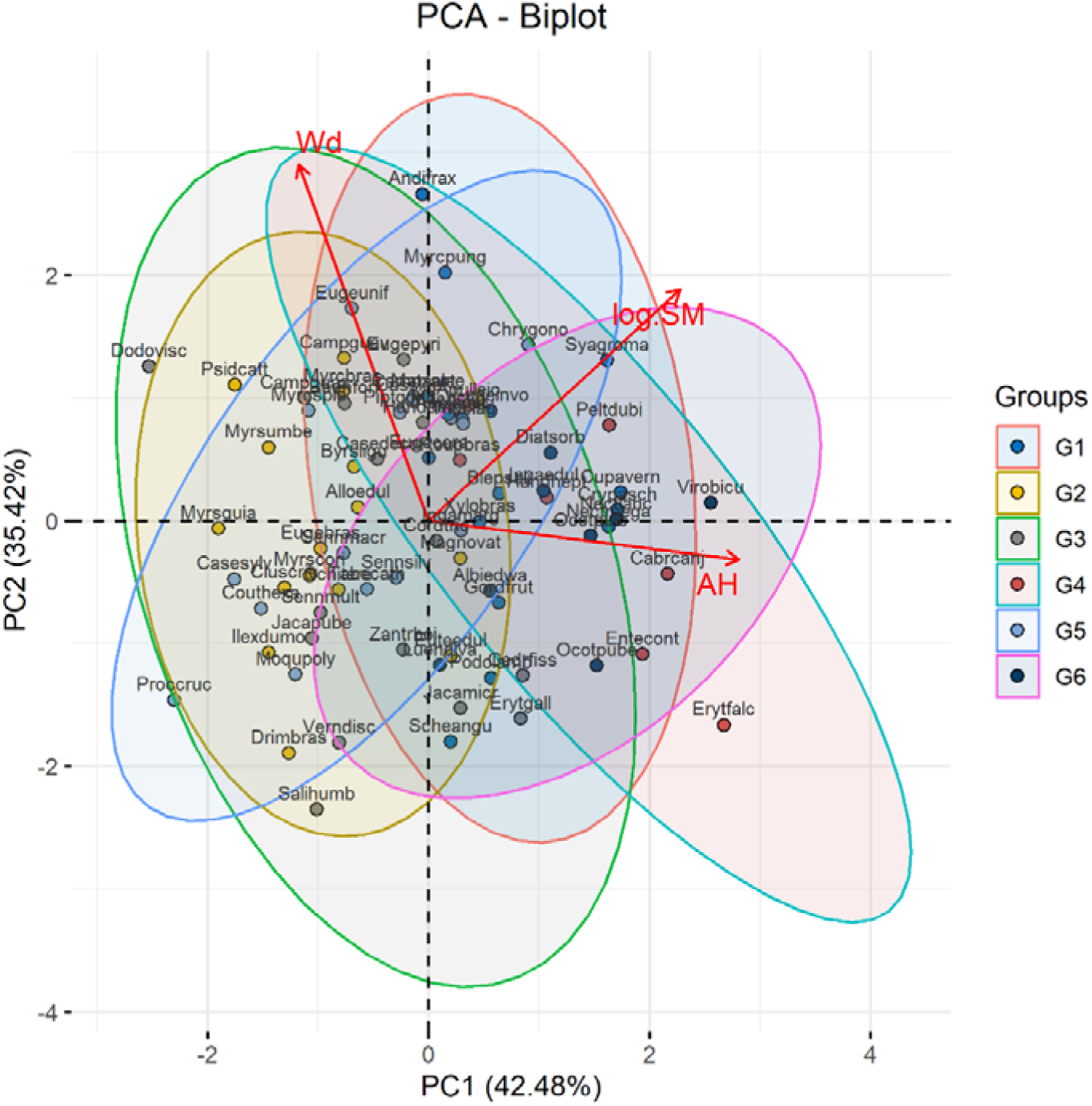
Principal component analysis (PCA) of the three continuous functional traits of the 77 species, representing the six functional groups from the cluster analysis. Species are represented with the same colors as the ellipse (functional group) to which they belong. The red lines and arrows represent the functional traits: wood density (Wd), seed mass (log.SM), and Average height (AH). The dotted lines indicate the main axes (PC 1-2). Ellipses represent a 95% confidence interval for a given functional group. For species codes and a complete list, see Supplementary material – Table S1.

## 4. Discussion

This is the first study that uses the functional group approach to conceive a tool for optimizing the diversification of the urban forest using native trees of the Atlantic Forest domain. The Atlantic Forest biome is classified as a priority area for conservation due to its significant diversity, endemism, and number of threatened species (Campanili and Schäffer, 2010). The state of Santa Catarina is completely inserted within the Atlantic Forest Biome and has a high richness of native species in its different phytophysiognomies, but this diversity is rarely used in urban environments. On top of improving the quality of ecological corridors connecting natural areas through urbanized areas, the use of native species in urban forests also has an important cultural role in the cities. The preservation of native species is important for retaining the biological distinctiveness of urban areas, but also for conservation goals. Promoting native biodiversity in urban areas not only educates city dwellers about local species but also enhances public awareness and emotional connection crucial for conservation efforts (McKinney, 2006).

From data of 77 native species from the Dense Ombrophilous Forest in Santa Catarina state, Brazil, we found six distinct functional groups (Fig. 2, Fig. 3) based on the functional traits analyzed (seed mass, wood density, average height, and leaf persistence). The different groups represent species with different ecological strategies. In group 1, we find persistent species that are more conservative in their use of water (Mediavilla and Escudero, 2003) and that were reported to have low photosynthetic capacity (Wright et al., 2006). The species in this group have moderate to slow growth and generally high wood density, which is associated with a decline in wood water content and, hence, its potential for water storage (Borchert, 1994), and are less able to exploit temporally favorable growth conditions (Rüger et al., 2012). High wood density also offers greater mechanical support against physical stresses such as winds and stores more carbon (Chave et al., 2009). The heavy seeds present in this group are related to greater resistance to environmental hazards due to larger reserves that can be exploited as a mechanism of drought resistance (Leishman and Westoby, 1994).

In functional group 2, we also have persistent species (more conservative in their use of water) that are also important for shade and for the conservation of biodiversity by providing foliage year-round (Núñez-Florez et al., 2019). Small-statured species with high wood density normally grow slowly (Rüger et al., 2012). They have greater mechanical support against physical stresses such as winds and store more carbon (Chave et al., 2009). The lighter seeds in this group characterize larger dispersal distances (Rüger et al., 2012; Muller-Landau et al., 2008), which is essential to the species’ survival and contributes to the seed dispersion in urban remnants of native forests.

In functional group 3, we have deciduous species that normally have a high rate of photosynthesis per unit leaf mass, low leaf longevity, and low specific leaf mass, which characterizes drought avoidance and tolerance strategies (Souza et al.; 2015). Deciduous species are also important in subtropical climates to allow more light availability in the urban environment during the winter season. This group is represented by pioneer and secondary species with fast to moderate growth. The light seeds in this group are an efficient dispersal strategy (Rüger et al., 2012; Muller-Landau et al., 2008).

Functional group 4 is represented by deciduous secondary species that express adaptation to seasonality and drought, resulting in reduced activities during the unfavorable season and resumption of growth with variable rates of resource use during the short favorable season (Singh and Singh, 1992). The species in this group also have moderate growth but high stature, which increases dispersal distances (Thomson et al., 2018) in combination with light seeds, which contributes to seed dispersion in urban remnants of native forests.

Functional group 5 is formed by semi-deciduous (tolerance strategy) pioneer and secondary species with short to medium stature, moderate growth, and heavy seeds, that are more resistant to environmental hazards (Leishman and Westoby, 1994).

Functional group 6 is represented by semi-deciduous (tolerance strategy), secondary species with high stature that respond more strongly to higher light (Rüger et al., 2012). Species in this group have fast to moderate growth, and heavy seeds, which characterize a greater resistance to environmental hazards (Leishman and Westoby, 1994).

To create more resistant urban forests in cities in the domain of DOF it is important to combine species from different groups. Greatest distances, and thus benefits, are found among species from different groups, which only become useful once maximum group diversity is achieved. Users can select species from various groups that align with site-specific conditions, optimizing functional diversity. This would increase the probability that the selected trees will maintain the flow of services when facing multiple and largely unpredictable stressors (Paquette et al., 2021). For example, we could combine *Blepharocalyx salicifolius* (1), *Eugenia pyriformis* (3), *Casearia Sylvestris* (5), *Clusia criuva* (2), *Peltophorum dubium* (4), and *Luehea divaricata* (6), that belong to different functional groups, that have different functional traits and consequently have different ecological strategies. Combining this high functional diversity will help the urban forest be more resistant to varying hazards from climate change. It is important to understand that we propose a tool to diversify the urban forest, not as a species selection tool; it is still important to pay attention to other characteristics of the species to determine if it is suitable for the location considered, given local restrictions and environmental characteristics (Núñez-Florez et al., 2019).

Similar projects have created functional groups to be used by managers by cluster analysis to diversify species in Urban Forests using functional traits (Paquette et al., 2021, Núñez-Florez et al., 2019). Both found an efficient and easy method to differentiate species and help managers plan more resilient forests.

We recognized that this study has limitations, especially related to the lack of information on functional traits for several species of DOF. Some important functional traits, such as leaf nitrogen content and specific leaf area, often used in ecology (Díaz et al., 2016; Reich 2014; Chave et al., 2009; Wright et al. 2004), including urban forests (Paquette et al., 2021), could not be used because of insufficient data. This reinforces the need for further research on functional traits of DOF species, and the global south in general.

Many of the species suggested are common in urban tree planting in the state, such as *Syagrus romanzoffiana*, *Handroanthus chrysotrichus*, *Handroanthus heptaphyllus*, *Peltophorum dubium*, among others, whose adaptation to urban environments has already been confirmed. These species are easily found in local nurseries and garden centers. Other species in our pool were selected based on their high potential for use in urban environments, but they are still little known, both in terms of their adaptation to urban environments and their availability. We hope that this research will encourage the use and production of these species in urban environments and that it will also generate new research into the use of native species in urban environments.

## 5. Conclusion

We used the functional group approach to design a tool to assist municipalities in increasing the diversity and resilience of the urban forest with the use of native trees of the Atlantic Forest domain. We classified 77 native trees of the Dense Ombrophilous Forest (DOF) in Santa Catarina state, southern Brazil, with potential use in urban environments into six functional groups according to their functional traits.

Using this approach will help create more resistant urban forests in cities in the domain of DOF by alternating species of different functional groups with different ecological functions, reducing the risk associated with global changes, such as increased environmental stress and pests. This method will also help incentivize the use of native species from DOF, which will improve the knowledge and connection of people with native trees, advance the benefits derived from the ecosystem services of provisioning, regulation, and culture, and help to promote their conservation. The same method could also be easily adapted to different biomes in Brazil and abroad.

## References

Abreu-Harbich, L.V. de, Labaki, L.C., Matzarakis, A., 2015. Effect of tree planting design and tree species on human thermal comfort in the tropics. Landscape and Urban Planning 138, 99–109. 10.1016/j.landurbplan.2015.02.008

Alvares, C.A., Stape, J.L., Sentelhas, P.C., Gonçalves, J.L.M., Sparovek, G., 2013. Köppen’s climate classification map for Brazil. Meteorologische zeitschrift 22(6), 711–728. 10.1127/0941-2948/2013/0507

Alves, L.P., Costa, J.A.S., Costa, C.B.N., 2023. Arborização urbana dominada por espécies exóticas em um país megadiverso: falta de planejamento ou desconhecimento? Revista Brasileira de Geografia Física 16(3), 1304–1375. (In Portuguese, with English summary). 10.26848/rbgf.v16.3.p1304-1375

Bechara, F.C., Topanotti, L.R., Silva, L. M., 2016. Aspectos da arborização urbana ecológica. Revista Ibero-Americana de Ciências Ambientais 7 (1), 49–55. (In Portuguese, with English summary). 10.6008/SPC2179-6858.2016.001.0004

Bernhardt-Römermann, M., Römermann, C., Nuske, R., Parth, A., Klotz, S., Schmidt, W., & Stadler, J., 2008. On the identification of the most suitable traits for plant functional trait analyses. Oikos 117(10), 1533–1541. 10.1111/j.0030-1299.2008.16776.x

Borchert, R., 1994. Soil and Stem Water Storage Determine Phenology and Distribution of Tropical Dry Forest Trees. Ecology 75(5), 1437–1449. 10.2307/1937467

Bush, J., Doyon, A., 2019. Building urban resilience with nature-based solutions: How can urban planning contribute? Cities 95, 102483. 10.1016/j.cities.2019.102483

Cadotte, M.W., Carscadden, K., Mirotchnick, N., 2011. Beyond species: functional diversity and the maintenance of ecological processes and services. Journal of Applied Ecology 48, 1079–1087. 10.1111/j.1365-2664.2011.02048.x

Campanili M, Schäffer WB., 2010. Mata Atlântica: patrimônio nacional dos brasileiros. Ministério do Meio Ambiente, Brasília. (In Portuguese)

Carvalho, P.E.R. 2003. Espécies Arbóreas Brasileiras. Embrapa, Vol. 1, Brasília. (In Portuguese)

Carvalho, P.E.R. 2006. Espécies Arbóreas Brasileiras. Embrapa, Vol. 2, Brasília. (In Portuguese)

Carvalho, P.E.R. 2008. Espécies Arbóreas Brasileiras. Embrapa, Vol. 3, Brasília. (In Portuguese)

Carvalho, P.E.R. 2010. Espécies Arbóreas Brasileiras. Embrapa, Vol. 4, Brasília. (In Portuguese)

Carvalho, P.E.R. 2014. Espécies Arbóreas Brasileiras. Embrapa, Vol. 5, Brasília. (In Portuguese)

Chave, J., Andalo, C., Brown, S., Cairns, M. A., Chambers, J.Q., Eamus, D., Folster, H., Fromard, F., Higuchi, N., Kira, T., Lescure, J.-P., Nelson, B.W., Ogawa, H., Puig, H., Riéra, B., Yamakura, T., 2005. Tree allometry and improved estimation of carbon stocks and balance in tropical forests. Oecologia 145, 87–99. 10.1007/s00442-005-0100-x

Chave, J., Coomes, D., Jansen, S., Lewis, S.L., Swenson, N.G., Zanne, A.E., 2009. Towards a worldwide wood economics spectrum. Ecology Letters 12, 351–66. 10.1111/j.1461-0248.2009.01285.x

Díaz, S., Kattge, J., Cornelissen, J.H.C., Wright, I.J., Lavorel, S., Dray, S., Reu, B., Kleyer, M., Wirth, C., Colin Prentice, I., Garnier, E., Bonisch, ^..^ G., Westoby, M., Poorter, H., Reich, P.B., Moles, A.T., Dickie, J., Gillison, A.N., Zanne, A.E., Chave, J., Joseph Wright, S., Sheremet Ev, S.N., Jactel, H., Baraloto, C., Cerabolini, B., Pierce, S., Shipley, B., Kirkup, D., Casanoves, F., Joswig, J.S., Günther, A., Falczuk, V., Rüger, N., Mahecha, M.D., Gorné, L.D., 2016. The global spectrum of plant form and function. Nature 529, 167–171. 10.1038/nature16489

Elias, G.A., Citadini-Zanette, V., Santos, R., 2020. Árvores nativas para a arborização urbana: um estudo de caso no Sul do Brasil. Revista Brasileira de Agroecologia 15(5), 249–260. (In Portuguese, with English summary). 10.33240/rba.v15i5.22907

Esperon-Rodriguez, M., Rymer, P.D., Power, S.A., Barton, D.N., Cariñanos, P., Dobbs, C., Eleuterio, A.A., Escobedo, F.J., Hauer, R., Hermy, M., Jahani, A., Onyekwelu, J.C., Östberg, J., Pataki, D., Randrup, T.B., Rasmussen, T., Roman, L. A., Russo, A., Shackleton, C., Solfjeld, I., van Doorn, N.S., Wells, M.J., Wiström, B., Yan, P., Yang, J., Tjoelker, M.G., 2022. Assessing climate risk to support urban forests in a changing climate. Plants, People, Planet 4(3), 201–213. 10.1002/ppp3.10240

Farrell, C., Livesley, S., Arndt, S., Beaumont, L., Burley, H., Ellsworth, D., Esperon-Rodriguez, M., Fletcher, T., Gallagher, R., Ossola, A., Power, S., Marchin, R., Rayner, J., Rymer, P., Staas, L., Szota, C., Williams, N., Leishman, M. 2022. Can we integrate ecological approaches to improve plant selection for green infrastructure? Urban Forestry & Urban Greening 76, 127732. 10.1016/j.ufug.2022.127732

Flora e Funga do Brasil 2020. Jardim Botânico do Rio de Janeiro. (In Portuguese). Retrieved December 12, 2023 from: http://floradobrasil.jbrj.gov.br/.

Garnier, E., Navas, M.L., 2012. A trait-based approach to comparative functional plant ecology: concepts, methods, and applications for agroecology. A review. Agronomy for Sustainable Development 32, 365–399. 10.1007/s13593-011-0036-y

Gasper, A. L., Uhlmann, A., Sevegnani, L., Meyer, L., Lingner, D.V., Verdi, M., Stival-Santos, A., Sobral, M., Vibrans, A.C., 2014. Floristic and Forest Inventory of Santa Catarina: species of evergreen rainforest. Rodriguésia 65(4), 807–816. 10.1590/2175-7860201465401

Gonçalves, L.M., Monteiro, P.H.S., Santos, L.S., Maia, N.J.C., Rosal, L.F. 2018. Arborização Urbana: a importância do seu planejamento para qualidade de vidas nas cidades. Ensaios e Ciência 22(2), 128–136. (In Portuguese, with English summary). 10.17921/1415-6938.2018v22n2p128-136

Gonçalves, W., 2009. Arborização urbana. Centro de Produções Técnicas, Viçosa, MG. (In Portuguese).

Guo, J., Xu, Q., Zeng, Y., Liu, Z., Zhu, X.X., 2023. Nationwide urban tree canopy mapping and coverage assessment in Brazil from high-resolution remote sensing images using deep learning, ISPRS Journal of Photogrammetry and Remote Sensing 198, 1–15. 10.1016/j.isprsjprs.2023.02.007

Harel, D., Holzapfel, C., Sternberg, M., 2011. Seed mass and dormancy of annual plant populations and communities decreases with aridity and rainfall predictability. Basic and Applied Ecology 12(8), 674–684. 10.1016/j.baae.2011.09.003

Husson, F., Josse, J., Le, S., Mazet, J., 2017. FactoMineR: Multivariate Exploratory Data Analysis and Data Mining. R package version 2.1.6. Retrieved December 12, 2023 from: http://cran.nexr.com/web/packages/FactoMineR/FactoMineR.pdf.

Isbell, F., Craven, D., Connolly, J., Loreau, M., Schmid, B., Beierkuhnlein, C., Bezemer, T.M., Bonin, C., Bruelheide, H., de Luca, E., Ebeling, A., Griffin, J.N., Guo, Q., Hautier, Y., Hector, A., Jentsch, A., Kreyling, J., Lanta, V., Manning, P., Meyer, S.T., Mori, A.S., Naeem, S., Niklaus, P.A., Polley, H.W., Reich, P.B., Roscher, C., Seabloom, E.W., Smith, M.D., Thakur, M.P., Tilman, D., Tracy, B.F., van der Putten, W.H., van Ruijven, J., Weigelt, A., Weisser, W.W., Wilsey, B., Eisenhauer, N., 2015. Biodiversity increases the resistance of ecosystem productivity to climate extremes. Nature, 526, 574–577. 10.1038/nature15374

Kassambara, A., Mundt, F., 2020. Factoextra: Extract and Visualize the Results of Multivariate Data Analyses. R Package Version 1.0.7. Retrieved December 12, 2023 from: https://CRAN.R-project.org/package=factoextra.

Kattge, J., Bönisch, G., Díaz, S., et al., 2020. TRY plant trait database – enhanced coverage and open access. Global Change Biology 26, 119–188. 10.1111/gcb.14904

Lachenbruch, B., McCulloh K.A., 2014. Traits, properties, and performance: how woody plants combine hydraulic and mechanical functions in a cell, tissue, or whole plant. New Phytologist 204, 747–764. 10.1111/nph.13035

Leishman, M.R., Westoby, M., 1994. The role of large seeds in seedling establishment in dry soil conditions - experimental evidence from semi-arid species. Journal of Ecology 82, 249–258. 10.2307/2261293

Lingner, D.V., Schorn, L.A., Vibrans, A.C., Meyer, L., Sevegnani, L., Gasper, A.L., Sobral, M.G., Krüger, A., Klemz, G., Schmidt, R., Anastácio Junior, C., 2013. Fitossociologia do componente arbóreo/arbustivo da Floresta Ombrófila Densa em Santa Catarina. In: Vibrans, A.C., Sevegnani, L., Gasper, A.L., Lingner, D.V. (Eds.). Inventário Florístico Florestal de Santa Catarina Volume IV – Floresta Ombrófila Densa. Edifurb, Blumenau, pp. 159–202. (In Portuguese)

Lorenzi, H., 1992. Árvores brasileiras: manual de identificação e cultivo de plantas arbóreas nativas do Brasil. Instituto Plantarum, Vol 1, Nova Odessa, SP. (In Portuguese).

Lorenzi, H., 1998. Árvores brasileiras: manual de identificação e cultivo de plantas arbóreas nativas do Brasil. Instituto Plantarum, Vol 2, Nova Odessa, SP. (In Portuguese).

Lorenzi, H., 2009. Árvores brasileiras: manual de identificação e cultivo de plantas arbóreas nativas do Brasil. Instituto Plantarum, Vol 3, Nova Odessa, SP. (In Portuguese).

Maechler, M., Rousseeuw, P., Struyf, A., Hubert, M., Hornik, K., 2023. cluster: cluster analysis basics and extensions. R package version 2.1.6. Retrieved December 12, 2023 from: https://CRAN.R-project.org/package=cluster

McKinney, M.L., 2006. Urbanization as a major cause of biotic homogenization. Biological Conservation 127(3), 247–260. 10.1016/j.biocon.2005.09.005

Mediavilla, S., Escudero, A., 2003. Photosynthetic capacity, integrated over the lifetime of a leaf, is predicted to be independent of leaf longevity in some tree species. New Phytologist 159(1), 203–211. 10.1046/j.1469-8137.2003.00798.x

Moles, A.T., Westoby, M., 2006. Seed size and plant strategy across the whole life cycle. Oikos 113, 91–105. 10.1111/j.0030-1299.2006.14194.x

Monalisa-Francisco, N., Ramos, F.N., 2019. Composition and Functional Diversity of the Urban Flora of Alfenas-MG, Brazil. Floresta e Ambiente 26(3), e20171110. 10.1590/2179-8087.111017

Moro, M.F., Castro, A.S.F., 2015. A checklist of plant species in the urban forestry of Fortaleza, Brazil: where are the native species in the country of megadiversity? Urban Ecosystems 18, 47–71. 10.1007/s11252-014-0380-1

Muller-Landau, H. C., S. J. Wright, O. Caldero n, R. Condit, and S. P. Hubbell., 2008. Interspecific variation in primary seed dispersal in a tropical forest. Journal of Ecology 96, 653–667. 10.1111/j.1365-2745.2008.01399.x

Myers, N., Mittermeier, R.A., Mittermeier, C.G., Fonseca, G.A.B., Kent, J., 2000. Biodiversity hotspots for conservation priorities. Nature 403, 853–845. 10.1038/35002501

Nock, C.A., Paquette, A., Follett, M., Nowak, D.J., Messier, C., 2013. Effects of urbanization on tree species functional diversity in Eastern North America. Ecosystems 16, 1487–1497. 10.1007/s10021-013-9797-5.

Núñez-Florez, R., Pérez-Gómez, U., Fernández-Méndez, F., 2019. Functional diversity criteria for selecting urban trees. Urban Forestry & Urban Greening 38, 251–266. 10.1016/j.ufug.2019.01.005.

Oksanen, F.J., Simpson, G.L., Blanchet, F. G., Kindt, R., Legendre, P., Minchin. P.R., O’Hara, R.B., Solymos, P., Stevens, M.H.H., Szoecs, E., Wagner, H., Barbour. M., Bedward, M., Bolker, B., Borcard, D., Carvalho, G., Chirico, M., Caceres, M. de, Durand, S., Evangelista, H.B.A., FitzJohn, R., Friendly, M., Furneaux, B., Hannigan, G., Hill, M.O., Lahti, L., McGlinn, D., Ouellette, M-H., Cunha, E.R., Smith, T., Stier, A., Ter Braak, C.J.F., Weedon, J., 2017. Vegan: Community Ecology Package. R package Version 2.4-3. Retrieved December 12, 2023 from: https://CRAN.R-project.org/package=vegan.

Oliveira-Filho, A.T., Budke, J.C., Jarenkow, J.A., Eisenlohr, P.V., Neves, D.R.M., 2015. Delving into the variations in tree species composition and richness across South American subtropical Atlantic and Pampean forests. Journal of Plant Ecology 8(3), 242–260. 10.1093/jpe/rtt058

Pamukcu-Albers, P., Ugolini, F., La Rosa, D., Grădinaru, S.R., Azevedo, J.C., Wu, J., 2021. Building green infrastructure to enhance urban resilience to climate change and pandemics. Landscape Ecology 36, 665–673. 10.1007/s10980-021-01212-y

Paquette, A., Sousa-Silva, R., Maure, F., Cameron, E., Belluau, M., Messier, C., 2021. Praise for diversity: A functional approach to reduce risks in urban forests. Urban Forestry & Urban Greening 62, 127157. 10.1016/j.ufug.2021.127157.

Pastorello, C.E. de S.P., Xavier, M.V.B., Fonseca, A.P.M., Aguiar, R.M.A.S., Figueiredo, M.A.P., 2022. Traços Funcionais e Estratos Verticais na Arborização de uma Praça de Montes Claros, Minas Gerais. Revista da Sociedade Brasileira de Arborização Urbana 17(4), 60–75. (In Portuguese, with English summary). 10.5380/revsbau.v17i4.87419

Pavoine, S., Vallet, J., Dufour, A.B., Gachet, S., Daniel, H., 2009. On the challenge of treating various types of variables: Application for improving the measurement of functional diversity. Oikos 118(3), 391–402. 10.1111/j.1600-0706.2008.16668.x

R Core Team, 2021. R: a language and environment for statistical computing. R Foundation for Statistical Computing, Vienna, Austria. Retrieved December 12, 2023 from: https://www.R-project.org/.

Reich, P.B., 2014. The world-wide ‘fast–slow’ plant economics spectrum: a traits manifesto. Journal of Ecology 102, 275–301. 10.1111/1365-2745.12211

Rezende, C.L., Scarano, F.R., Assad, E.D., Joly, C.A., Metzger, J.P., Strassburg, B.B.N., Tabarelli, M., Fonseca, G.A., Mittermeier, R.A., 2018. From hotspot to hopespot: An opportunity for the Brazilian Atlantic Forest. Perspectives in Ecology and Conservation 16(4), 208–214. 10.1016/j.pecon.2018.10.002

Rüger, N., Wirth, C., Wright, S.J., Condit, R., 2012. Functional traits explain light and size response of growth rates in tropical tree species. Ecology 93(12), 2626–2636. 10.1890/12-0622.1

Sartori, R.A., Martins, G.A., Zaú, A.S., Brasil, L.S.C., 2019. Urban afforestation and favela: A study in A community of Rio de Janeiro, Brazil. Urban Forestry & Urban Greening 40, p. 84–92. 10.1016/j.ufug.2018.10.004

Saueressig D., 2014. Plantas do Brasil: Árvores Nativas. Editora Plantas do Brasil, Vol 1, Irati, PR. (In Portuguese).

Seddon, N., Chausson, A., Berry, P., Girardin, C.A.J., Smith, A., Turner, B., 2020. Understanding the value and limits of nature-based solutions to climate change and other global challenges. Philosophical Transactions of the Royal Society B 375(1794), 20190120. 10.1098/rstb.2019.0120

Sevegnani, L., Gasper, A.L., Bonnet, A., Sobral, M.G., Vibrans, A.C., Verdi, M., Santos, A.S., Dreveck, S., Korte, A., Schmitt, J., Cadorin, T., Lopes, C.P., Cagliono, E., Torres, J.F., Meyer, L., 2013. Flora Vascular da Floresta Ombrófila Densa em Santa Catarina. In: Vibrans, A.C., Sevegnani, L., Gasper, A.L., Lingner, D.V. (Eds.). Inventário Florístico Florestal de Santa Catarina Vol. IV - Floresta Ombrófila Densa. Edifurb, Blumenau, pp. 127–142. (In Portuguese)

SFB - Serviço Florestal Brasileiro. Inventário florestal nacional: principais resultados: Santa Catarina. Brasília, DF, 2018. Retrieved October 25, 2024 from: https://snif.florestal.gov.br/images/pdf/publicacoes/periodo_eleitoral/publicacoes_ifn/relatorios/IFN_SC_2018_periodo_eleitoral.pdf

Singh, J.S., Singh, V.K., 1992. Phenology of seasonally dry tropical forest. Current Science 63, 684–689. Retrieved January 8, 2024 from: https://www.jstor.org/stable/24094777

Soares, A.C.S., Santos, R.O. dos, Soares, R.N., Cantuaria, P.C., de Lima, R.B. de, Breno Silva e Silva, B.M. da., 2021. Paradox of afforestation in cities in the Brazilian Amazon: An understanding of the composition and floristic similarity of these urban green spaces. Urban Forestry & Urban Greening 66, 127374. 10.1016/j.ufug.2021.127374

Souza, B.C. de, Oliveira, R.S., Araújo, F.S. de, Lima, A.L.A. de, Rodal, M.J.N., 2015. Divergências funcionais e estratégias de resistência à seca entre espécies decíduas e sempre verdes tropicais. Rodriguésia 66(1), 21–32. (In Portuguese, with English summary). 10.1590/2175-7860201566102

Tilman, D., 1999. The ecological consequences of changes in biodiversity: A search for general principles. Ecology 80, 1455–1474. 10.1890/0012-9658(1999)080[1455:TECOCI]2.0.CO,2

Thomson, F.J., Letten, A.D., Tamme, R., Edwards, W. and Moles, A.T., 2018. Can dispersal investment explain why tall plant species achieve longer dispersal distances than short plant species? New Phytologist 217, 407–415. 10.1111/nph.14735

USDA - U.S. Department of Agriculture Forest Service, 2006. The Urban Forestry Manual: A Manual for Urban Forestry in the South. U.S. Department of Agriculture Forest Service Southern Research Station, Athens, GA. Forests, Management, Manual/guidance document, Urban, Forestry. Retrieved January 8, 2024 from: https://www.fao.org/sustainable-forest-management/toolbox/tools/tool-detail/en/c/284850/

Vargas-Hernández, J.G., Pallagst, K., Zdunek-Wielgołaska, J., 2018. Urban Green Spaces as a Component of an Ecosystem. In: Marques, J. (Ed.). Handbook of Engaged Sustainability. Springer, Cham., pp. 885–916. 10.1007/978-3-319-71312-0_49

Vibrans, A.C., Nicoletti, A.L., Liesenberg, V., Refosco, J.C., Kohler, L.P.A., Bizon, A. R., Lingner, D.V., Bosco, F.D., Bueno, M.M., Silva, M.S., Pessatti, T.B., 2021. Monitora SC: um novo mapa de cobertura florestal e uso da terra do estado de Santa Catarina. Agropecuária Catarinense 34(2), 42-. 10.52945/rac.v34i2.1086

Wang, Z., Li, Y., Su, X., Tao, S., Feng, X., Wang, Q., Xu, X., Liu, Y., Michaletz, S.T., Shrestha, N., Larjavaara, M., Enquist, B.J., 2019. Patterns and ecological determinants of woody plant height in eastern Eurasia and its relation to primary productivity. Journal of Plant Ecology 12(5), 791–803. 10.1093/jpe/rtz025

Westoby, M., Leishman, M., Lord, J., 1996. Comparative ecology of seed size and dispersal. Biological Sciences 351(1345), 1309–1318. 10.1098/rstb.1996.0114

Wright, I.J., Leishman, M.R., Read, C., Westoby, M., 2006. Gradients of light availability and leaf traits with leaf age and canopy position in 28 Australian shrubs and trees. Functional Plant Biology 33, 407–419. 10.1071/FP05319

Wright, I.J., Reich, P.B., Westoby, M., Ackerly, D.D., Baruch, Z., Bongers, F., Cavender-Bares, J., Chapin, T., Cornelissen, J.H.C., Diemer, M., Flexas, J., Garnier, E., Groom, P.K., Gulias, J., Hikosaka, K., Lamont, B.B., Lee, T., Lee, W., Lusk, C., Midgley, J.J., Navas, M-L., Niinemets, Ü., Oleksyn, J., Osada, N., Poorter, H., Poot, P., Prior, L., Pyankov, V.I., Roumet, C., Thomas, S.C., Tjoelker, M.G., Veneklaas, E.J., Villar, R., 2004. The worldwide leaf economics spectrum. Nature 428, 821–7. 10.1038/nature02403

Yamane, L., Botsford, L.W., Kilduff, D.P., 2018. Tracking restoration of population diversity via the portfolio effect. Journal of Applied Ecology 55(2), 472–481. 10.1111/1365-2664.12978

Zanne, A.E., Lopez-Gonzalez, G., Coomes, D. A., Ilic, J., Jansen, S., Lewis, S.L., Miller, R.B., Swenson, N.G., Wiemann, M.C., Chave, J., 2009. Data from: Towards a worldwide wood economics spectrum [Dataset]. Dryad. 10.5061/dryad.234

Zappi, D.C., Lovo, J., Hiura, A., Andrino, C.O., Barbosa-Silva, R.G., Martello, F., Gadelha-Silva, L., Viana, P.L., Giannini, T.C., 2022. Telling the Wood from the Trees: Ranking a Tree Species List to Aid Urban Afforestation in the Amazon. Sustainability 14, 1321. 10.3390/su14031321

Zhang, B., Brack, C.L., 2021. Urban forest responses to climate change: A case study in Canberra. Urban Forestry & Urban Greening 57, 126910. 10.1016/j.ufug.2020.126910

