## Supplemental Table S1 for "Native species of the Atlantic Forest for urban environments based on functional groups: an approach to make cities in southern Brazil more resilient"

**Table S1.** List of the 100 initial native species (species, genus, and family) of Dense Ombrophilous Forest with potential to use in the urban environment.

| **Species** | **Genus** | **Family** |
| --- | --- | --- |
| *Schinus terebinthifolia* | *Schinus* | Anacardiaceae |
| *Xylopia brasiliensis* | *Xylopia* | Annonaceae |
| *Annona rugulosa* | *Annona* | Annonaceae |
| *Annona sylvatica* | *Annona* | Annonaceae |
| *Aspidosperma australe* | *Aspidosperma* | Apocynaceae |
| *Tabernaemontana catharinensis* | *Tabernaemontana* | Apocynaceae |
| *Ilex dumosa* | *Ilex* | Aquifoliaceae |
| *Ilex brevicuspis* | *Ilex* | Aquifoliaceae |
| *Schefflera angustissima* | *Schefflera* | Araliaceae |
| *Euterpe edulis* | *Euterpe* | Arecaceae |
| *Syagrus romanzoffiana* | *Syagrus* | Arecaceae |
| *Moquiniastrum polymorphum* | *Moquiniastrum* | Asteraceae |
| *Vernonanthura discolor* | *Vernonanthura* | Asteraceae |
| *Handroanthus chrysotrichus* | *Handroanthus* | Bignoniaceae |
| *Handroanthus heptaphyllus* | *Handroanthus* | Bignoniaceae |
| *Jacaranda micrantha* | *Jacaranda* | Bignoniaceae |
| *Jacaranda puberula* | *Jacaranda* | Bignoniaceae |
| *Handroanthus umbellatus* | *Handroanthus* | Bignoniaceae |
| *Cordia trichotoma* | *Cordia* | Boraginaceae |
| *Clusia criuva* | *Clusia* | Clusiaceae |
| *Terminalia triflora* | *Terminalia* | Combretaceae |
| *Heisteria silvianii* | *Heisteria* | Erythropalaceae |
| *Albizia edwallii* | *Albizia* | Fabaceae |
| *Andira fraxinifolia* | *Andira* | Fabaceae |
| *Apuleia leiocarpa* | *Apuleia* | Fabaceae |
| *Bauhinia forficata* | *Bauhinia* | Fabaceae |
| *Cassia leptophylla* | *Cassia* | Fabaceae |
| *Enterolobium contortisiliquum* | *Enterolobium* | Fabaceae |
| *Erythrina crista-galli* | *Erythrina* | Fabaceae |
| *Erythrina falcata* | *Erythrina* | Fabaceae |
| *Inga edulis* | *Inga* | Fabaceae |
| *Inga marginata* | *Inga* | Fabaceae |
| *Machaerium stipitatum* | *Machaerium* | Fabaceae |
| *Peltophorum dubium* | *Peltophorum* | Fabaceae |
| *Piptadenia gonoachantha* | *Piptadenia* | Fabaceae |
| *Senna macranthera* | *Senna* | Fabaceae |
| *Senna multijuga* | *Senna* | Fabaceae |
| *Senna silvestris* | *Senna* | Fabaceae |
| *Andira anthelmia* | *Andira* | Fabaceae |
| *Ateleia glazioveana* | *Ateleia* | Fabaceae |
| *Dalbergia brasiliensis* | *Dalbergia* | Fabaceae |
| *Machaerium paraguariense* | *Machaerium* | Fabaceae |
| *Myrocarpus frondosus* | *Myrocarpus* | Fabaceae |
| *Vitex megapotamica* | *Vitex* | Lamiaceae |
| *Cryptocarya aschersoniana* | *Cryptocarya* | Lauraceae |
| *Nectandra lanceolata* | *Nectandra* | Lauraceae |
| *Nectandra megapotamica* | *Nectandra* | Lauraceae |
| *Ocotea puberula* | *Ocotea* | Lauraceae |
| *Ocotea pulchella* | *Ocotea* | Lauraceae |
| *Nectandra oppositifolia* | *Nectandra* | Lauraceae |
| *Cariniana estrellensis* | *Cariniana* | Lecythidaceae |
| *Magnolia ovata* | *Magnolia* | Magnoliaceae |
| *Byrsonima ligustrifolia* | *Byrsonima* | Malpighiaceae |
| *Luehea divaricata* | *Luehea* | Malvaceae |
| *Pleroma sellowianum* | *Pleroma* | Melastomataceae |
| *Tibouchina pulchra* | *Tibouchina* | Melastomataceae |
| *Cabralea canjerana* | *Cabralea* | Meliaceae |
| *Cedrela fissilis* | *Cedrela* | Meliaceae |
| *Trichilia clausseni* | *Trichilia* | Meliaceae |
| *Virola bicuhyba* | *Virola* | Myristicaceae |
| *Blepharocalyx salicifolius* | *Blepharocalyx* | Myrtaceae |
| *Campomanesia guaviroba* | *Campomanesia* | Myrtaceae |
| *Campomanesia guazumifolia* | *Campomanesia* | Myrtaceae |
| *Campomanesia xanthocarpa* | *Campomanesia* | Myrtaceae |
| *Eugenia brasiliensis* | *Eugenia* | Myrtaceae |
| *Eugenia cerasiflora* | *Eugenia* | Myrtaceae |
| *Eugenia involucrata* | *Eugenia* | Myrtaceae |
| *Eugenia pyriformis* | *Eugenia* | Myrtaceae |
| *Eugenia uniflora* | *Eugenia* | Myrtaceae |
| *Myrcia brasiliensis* | *Myrcia* | Myrtaceae |
| *Myrcia splendens* | *Myrcia* | Myrtaceae |
| *Myrcianthes pungens* | *Myrcianthes* | Myrtaceae |
| *Psidium cattleyanum* | *Psidium* | Myrtaceae |
| *Myrcianthes gigantea* | *Myrcianthes* | Myrtaceae |
| *Myrrhinium atropurpureum* | *Myrrhunium* | Myrtaceae |
| *Plinia peruviana* | *Plinia* | Myrtaceae |
| *Ouratea parviflora* | *Ouratea* | Ochnaceae |
| *Podocarpus lambertii* | *Podocarpus* | Podocarpaceae |
| *Myrsine coriacea* | *Myrsine* | Primulaceae |
| *Myrsine guianensis* | *Myrsine* | Primulaceae |
| *Myrsine umbellata* | *Myrsine* | Primulaceae |
| *Roupala montana* var*. brasiliensis* | *Roupala* | Proteaceae |
| *Coutarea hexandra* | *Coutarea* | Rubiaceae |
| *Zanthoxylum rhoifolium* | *Zanthoxylum* | Rutaceae |
| *Casearia decandra* | *Casearia* | Salicaceae |
| *Casearia sylvestris* | *Casearia* | Salicaceae |
| *Prockia crucis* | *Prockia* | Salicaceae |
| *Salix humboldtiana* | *Salix* | Salicaceae |
| *Allophylus edulis* | *Allophylus* | Sapindaceae |
| *Cupania vernalis* | *Cupania* | Sapindaceae |
| *Diatenopteryx sorbifolia* | *Diatenopteryx* | Sapindaceae |
| *Dodonaea viscosa* | *Dodonaea* | Sapindaceae |
| *Matayba elaeagnoides* | *Matayba* | Sapindaceae |
| *Chrysophyllum gonocarpum* | *Chrysophyllum* | Sapotaceae |
| *Brunfelsia uniflora* | *Brunfelsia* | Solanaceae |
| *Symplocos uniflora* | *Symplocos* | Symplocaceae |
| *Gordonia fruticosa* | *Gordonia* | Theaceae |
| *Citharexylum myrianthum* | *Cytharexylum* | Verbenaceae |
| *Duranta vestita* | *Duranta* | Verbenaceae |
| *Drimys brasiliensis* | *Drimys* | Winteraceae |
