## Supplemental Table S2 for "Native species of the Atlantic Forest for urban environments based on functional groups: an approach to make cities in southern Brazil more resilient"

**Table S2.** List of the 77 species (species, genus, and family) with their respective codes and functional groups.

| **Code** | **Species** | **Genus** | **Family** | **Group** |
| --- | --- | --- | --- | --- |
| Schitere | *Schinus terebinthifolia* Raddi | *Schinus* | Anacardiaceae | 1 |
| Xylobras | *Xylopia brasiliensis* Spreng | *Xylopia* | Annonaceae | 1 |
| Ilexdumo | *Ilex dumosa* Reissek | *Ilex* | Aquifoliaceae | 2 |
| Scheangu | *Schefflera angustissima* (Marchal) Frodin | *Schefflera* | Araliaceae | 1 |
| Syagroma | *Syagrus romanzoffiana* (Cham.) Glassman | *Syagrus* | Arecaceae | 1 |
| Euteedul | *Euterpe edulis* Mart. | *Euterpe* | Arecaceae | 2 |
| Cluscriu | *Clusia criuva* Cambess. | *Clusia* | Clusiaceae | 2 |
| Nectmega | *Nectandra megapotamica* (Spreng.) Mez | *Nectandra* | Lauraceae | 1 |
| Magnovat | *Magnolia ovata* (A.St.-Hil.) Spreng. | *Magnolia* | Magnoliaceae | 2 |
| Byrsligu | *Byrsonima ligustrifolia* A.Juss. | *Byrsonima* | Malpighiaceae | 2 |
| Eugeinvo | *Eugenia involucrata* DC. | *Eugenia* | Myrtaceae | 1 |
| Myrcbras | *Myrcia brasiliensis* Kiaersk. | *Myrcia* | Myrtaceae | 2 |
| Podolamb | *Podocarpus lambertii* Klotzsch ex Endl. | *Podocarpus* | Podocarpaceae | 1 |
| Myrscori | *Myrsine coriacea* (Sw.) R.Br. ex Roem. & Schult. | *Myrsine* | Primulaceae | 2 |
| Mataelae | *Matayba elaeagnoides* Radlk. | *Matayba* | Sapindaceae | 1 |
| Alloedul | *Allophylus edulis* (A.St.-Hil. et al.) Hieron. ex Niederl. | *Allophylus* | Sapindaceae | 2 |
| Gordfrut | *Gordonia fruticosa* (Schrad.) H.Keng | *Gordonia* | Theaceae | 1 |
| Drimbras | *Drimys brasiliensis* Miers | *Drimys* | Winteraceae | 2 |
| Myrcpung | *Myrcianthes pungens* (O.Berg) D.Legrand | *Myrcianthes* | Myrtaceae | 1 |
| Psidcatt | *Psidium cattleyanum* Sabine | *Psidium* | Myrtaceae | 2 |
| Heissilv | *Heisteria silvianii* Schwacke | *Heisteria* | Erythropalaceae | 1 |
| Andifrax | *Andira fraxinifolia* Benth. | *Andira* | Fabaceae | 1 |
| Crypasch | *Cryptocarya aschersoniana* Mez | *Cryptocarya* | Lauraceae | 1 |
| Campguav | *Campomanesia guaviroba* (DC.) Kiaersk. | *Campomanesia* | Myrtaceae | 2 |
| Eugecera | *Eugenia cerasiflora* Miq. | *Eugenia* | Myrtaceae | 1 |
| Blepsali | *Blepharocalyx salicifolius* (Kunth) O.Berg | *Blepharocalyx* | Myrtaceae | 1 |
| Eugebras | *Eugenia brasiliensis* Lam. | *Eugenia* | Myrtaceae | 2 |
| Myrsumbe | *Myrsine umbellata* Mart. | *Myrsine* | Primulaceae | 2 |
| Myrsguia | *Myrsine guianensis* (Aubl.) Kuntze | *Myrsine* | Primulaceae | 2 |
| Cupavern | *Cupania vernalis* Cambess. | *Cupania* | Sapindaceae | 1 |
| Machstip | *Machaerium stipitatum* Vogel | *Machaerium* | Fabaceae | 1 |
| Aspiaust | *Aspidosperma australe* Müll.Arg. | *Aspidosperma* | Apocynaceae | 5 |
| Verndisc | *Vernonanthura discolor* (Spreng.) H.Rob. | *Vernonanthura* | Asteraceae | 3 |
| Moqupoly | *Moquiniastrum polymorphum* (Less.) G. Sancho | *Moquiniastrum* | Asteraceae | 5 |
| Handchry | *Handroanthus chrysotrichus* (Mart. ex DC.) Mattos | *Handroanthus* | Bignoniaceae | 3 |
| Jacapube | *Jacaranda puberula* Cham. | *Jacaranda* | Bignoniaceae | 3 |
| Erytcris | *Erythrina crista-galli* L. | *Erythrina* | Fabaceae | 3 |
| Apulleio | *Apuleia leiocarpa* (Vogel) J.F.Macbr. | *Apuleia* | Fabaceae | 4 |
| Piptgono | *Piptadenia gonoacantha* (Mart.) J.F.Macbr. | *Piptadenia* | Fabaceae | 5 |
| Nectlanc | *Nectandra lanceolata* Nees | *Nectandra* | Lauraceae | 6 |
| Ocotpube | *Ocotea puberula* (Rich.) Nees | *Ocotea* | Lauraceae | 6 |
| Ocotpulc | *Ocotea pulchella* (Nees & Mart.) Mez | *Ocotea* | Lauraceae | 6 |
| Cabrcanj | *Cabralea canjerana* (Vell.) Mart. | *Cabralea* | Meliaceae | 3 |
| Virobicu | *Virola bicuhyba* (Schott ex Spreng.) Warb. | *Virola* | Myristicaceae | 6 |
| Couthexa | *Coutarea hexandra* (Jacq.) K.Schum. | *Coutarea* | Rubiaceae | 5 |
| Casedeca | *Casearia decandra* Jacq. | *Casearia* | Salicaceae | 3 |
| Casesylv | *Casearia sylvestris* Sw. | *Casearia* | Salicaceae | 5 |
| Salihumb | *Salix humboldtiana* Willd. | *Salix* | Salicaceae | 3 |
| Proccruc | *Prockia crucis* P.Browne ex L. | *Prockia* | Salicaceae | 5 |
| Diatsorb | *Diatenopteryx sorbifolia* Radlk. | *Diatenopteryx* | Sapindaceae | 6 |
| Chrygono | *Chrysophyllum gonocarpum* (Mart. & Eichler ex Miq.) Engl*.* | *Chrysophyllum* | Sapotaceae | 5 |
| Albiedwa | *Albizia edwallii* (Hoehne) Barneby & J.W.Grimes | *Albizia* | Fabaceae | 6 |
| Campxant | *Campomanesia xanthocarpa* (Mart.) O.Berg | *Campomanesia* | Myrtaceae | 5 |
| Casslept | *Cassia leptophylla* Vogel | *Cassia* | Fabaceae | 5 |
| Cedrfiss | *Cedrela fissilis* Vell. | *Cedrela* | Meliaceae | 3 |
| Cordtric | *Cordia trichotoma (Vell.)* Arráb. ex Steud. | *Cordia* | Boraginaceae | 3 |
| Erytfalc | *Erythrina falcata* Benth. | *Erythrina* | Fabaceae | 4 |
| Eugepyri | *Eugenia pyriformis* Cambess. | *Eugenia* | Myrtaceae | 3 |
| Jacamicr | *Jacaranda micranta* Cham. | *Jacaranda* | Bignoniaceae | 3 |
| Luehdiva | *Luehea divaricata* Mart. | *Luehea* | Malvaceae | 6 |
| Ingaedul | *Inga edulis* Mart. | *Inga* | Fabaceae | 6 |
| Entecont | *Enterolobium contortisiliquum* (Vell.) Morong | *Enterolobium* | Fabaceae | 4 |
| Ingamarg | *Inga marginata* Willd. | *Inga* | Fabaceae | 5 |
| Peltdubi | *Peltophorum dubium* (Spreng.) Taub. | *Peltophorum* | Fabaceae | 4 |
| Bauhforf | *Bauhinia forficata* Link | *Bauhinia* | Fabaceae | 3 |
| Sennmacr | *Senna macranthera* (DC. ex Collad.) H.S.Irwin & Barneby | *Senna* | Fabaceae | 5 |
| Sennsilv | *Senna silvestres* (Vell.) H.S.Irwin & Barneby | *Senna* | Fabaceae | 5 |
| Sennmult | *Senna multijuga* (Rich.) H.S.Irwin & Barneby | *Senna* | Fabaceae | 3 |
| Eugeunif | *Eugenia uniflora* L. | *Eugenia* | Myrtaceae | 5 |
| Myrcsple | *Myrcia splendens* (Sw.) DC. | *Myrcia* | Myrtaceae | 5 |
| Roupbras | *Roupala montana var. brasiliensis* (Klotzsch) K.S.Edwards | *Roupala* | Proteaceae | 4 |
| Zantrhoi | *Zanthoxylum rhoifolium* Lam. | *Zanthoxylum* | Rutaceae | 3 |
| Dodovisc | *Dodonaea viscosa* Jacq. | *Dodonaea* | Sapindaceae | 3 |
| Campguaz | *Campomanesia guazumifolia* (Cambess.) O.Berg | *Campomanesia* | Myrtaceae | 3 |
| Handhept | *Handroanthus heptaphyllus* (Vell.) Mattos | *Handroanthus* | Bignoniaceae | 4 |
| Tabecath | *Tabernaemontana catharinensis* A.DC. | *Tabernaemontana* | Apocynaceae | 5 |
| Tricclau | *Trichilia clausseni* C.DC. | *Trichilia* | Meliaceae | 5 |
