## Supplemental Table S3 for "Native species of the Atlantic Forest for urban environments based on functional groups: an approach to make cities in southern Brazil more resilient"

**Table S3.** Loadings and contribution of variables (%) for each PCA axis. R-square values in bold indicate the highest correlations of the variables with the axes.

|  | Loading (R-square) | | Contribution (%) | |
| --- | --- | --- | --- | --- |
| Functional traits | PC1 | PC2 | PC1 | PC2 |
| log.SM | **0.676** | **0.560** | 35.85 | 29.48 |
| AH | **0.834** | - 0.094 | 54.59 | 0.82 |
| Wd | - 0.349 | **0.861** | 9.55 | 69.69 |
