## Supplemental Table S4 for "Native species of the Atlantic Forest for urban environments based on functional groups: an approach to make cities in southern Brazil more resilient"

**Table S4.** Mean values and standard deviations of functional traits for each of the six groups.

|  | Functional traits | | |
| --- | --- | --- | --- |
| Groups | AH | Wd | SM |
| G1 | 15.88 ± 4.05 | 0.669 ± 0.144 | 0.736 ± 1.777 |
| G2 | 7.47 ± 2.45 | 0.652 ± 0.140 | 0.054 ± 0.070 |
| G3 | 10.13 ± 4.41 | 0.605 ± 0.169 | 0.151 ± 0.269 |
| G4 | 21.86 ± 3.02 | 0.604 ± 0.192 | 0.181 ± 0.116 |
| G5 | 9.56 ± 3.60 | 0.679 ± 0.099 | 0.483 ± 1.400 |
| G6 | 20.25 ± 3.85 | 0.582 ± 0.089 | 0.322 ± 0.430 |
